# Pulmonary lesions following inoculation with the SARS-CoV-2 Omicron BA.1 (B.1.1.529) variant in Syrian golden hamsters

**DOI:** 10.1101/2022.03.15.484448

**Authors:** Melanie Rissmann, Danny Noack, Debby van Riel, Katharina S. Schmitz, Rory D. de Vries, Peter van Run, Mart M. Lamers, Corine H. GeurtsvanKessel, Marion P. G. Koopmans, Ron A. M. Fouchier, Thijs Kuiken, Bart L. Haagmans, Barry Rockx

## Abstract

The Omicron BA.1 (B.1.1.529) SARS-CoV-2 variant is characterized by a high number of mutations in the viral genome, associated with immune-escape and increased viral spread. It remains unclear whether milder COVID-19 disease progression observed after infection with Omicron BA.1 in humans is due to reduced pathogenicity of the virus or due to pre-existing immunity from vaccination or previous infection. Here, we inoculated hamsters with Omicron BA.1 to evaluate pathogenicity and kinetics of viral shedding, compared to Delta (B.1.617.2) and to animals re-challenged with Omicron BA.1 after previous SARS-CoV-2 614G infection. Omicron BA.1 infected animals showed reduced clinical signs, pathological changes, and viral shedding, compared to Delta-infected animals, but still showed gross- and histopathological evidence of pneumonia. Pre-existing immunity reduced viral shedding and protected against pneumonia. Our data indicate that the observed decrease of disease severity is in part due to intrinsic properties of the Omicron BA.1 variant.

## Background

On 31^st^ December 2019, the World Health Organization (WHO) was informed of a cluster of cases of pneumonia of unknown cause detected in Wuhan City, Hubei Province of China. A novel coronavirus (SARS-CoV-2) was identified [1]. Characterized by high transmissibility, this novel disease named coronavirus disease 2019 (COVID-19) rapidly spread throughout the whole world. As of March 9^th^ 2022, WHO has reported almost 440 million SARS-CoV-2 infections with over 6 million confirmed deaths worldwide [2]. The WHO has been monitoring and assessing the evolution of SARS-CoV-2 since its emergence and designated specific Variants of Concern (VOCs) that pose an increased risk to global public health. To date, several VOCs of SARS-CoV-2 have been identified with sets of mutations in the spike protein or other sites in the viral genome, linked to increased transmission or disease, or evasion of immune responses [3].

The VOC Omicron BA.1 (B.1.1.529) was initially reported in South Africa and Botswana in November 2021 [4], but has spread rapidly and is the predominant strain replacing the previously dominant Delta strain (B.1.617.2) in many countries worldwide. A striking feature of this variant is that it has an unusual large number of mutations in the spike protein, leading to partial evasion of pre-existing antibodies, and rendering it unsusceptible to the majority of monoclonal antibody therapies approved for clinical use [5–7]. In addition, epidemiological data suggest that infection with the Omicron BA.1 variant leads to less severe disease in humans [8], although it remains unknown whether this decrease in pathogenicity is primarily caused by intrinsic properties of the virus or if the observed milder course of disease in humans is due to pre-existing immunity, either from vaccination or previous exposure to other SARS-CoV-2 variants [9]. A variant dependent decrease of disease severity was supported by first observations of limited pathogenicity of Omicron BA.1 in immunologically naïve Syrian golden hamsters [10, 11].

To evaluate the intrinsic pathogenic potential of the emerging Omicron BA.1 variant, groups of Syrian golden hamsters were infected with Omicron BA.1. Results were compared to hamsters infected with the Delta variant. In addition, we evaluated whether previous exposure to the ancestral SARS-CoV-2 strain 614G affords protection against heterologous challenge with the antigenically distinct Omicron BA.1 variant.

## Methods

### Viruses and cells

Calu-3 cells were cultured in Opti-MEM I (1) + GlutaMAX (Gibco) supplemented with 10% FBS, penicillin (100 IU/mL), and streptomycin (100 IU/mL) at 37°C in a humidified CO_2_ incubator. Cells were tested negative for mycoplasma. Bavpat-1 (614G), Delta (B.1.617.2) and Omicron BA.1 (B.1.1.529) variants of SARS-CoV-2 were propagated as described before [12].Briefly, all used viruses were propagated to passage 3 on Calu-3 (ATCC HTB-55) cells in Advanced DMEM/F12 (Gibco), supplemented with HEPES, Glutamax, penicillin (100 IU/mL) and streptomycin (100 IU/mL) (AdDF+++) at 37°C in a humidified CO2 incubator. Infections were performed at a multiplicity of infection (MOI) of 0.01 and virus was harvested after 72 hours. The culture supernatant was cleared by centrifugation at 1000 × g for 5 min and stored at −80°C in aliquots. Virus titers were determined by plaque assay as described below. The Delta and Omicron BA.1 sequences are available on GenBank under accession numbers OM287123 and OM287553, respectively. All work with infectious SARS-CoV-2 was performed in a Class II Biosafety Cabinet under BSL-3 conditions at Erasmus Medical Center.

### Animals and experimental setup

Female Syrian golden hamsters (*Mesocricetus auratus*; 6 weeks old; Janvier, France) were handled in an ABSL-3 biocontainment laboratory (Supplementary material). Groups of animals (*n* = 8) were inoculated intranasally with 5 x 10^4^ PFU of the Omicron BA.1 or Delta variants of SARS-CoV-2 or PBS (mock; *n* = 4) in a total volume of 100 μl per animal. Throat swabs and nasal washes (250 μL) were taken at 1, 3, 5, 7, 10, 14, 18 and 21 days post inoculation (dpi). The body weight of animals was measured daily for the first 10 days of the experiment and at all days of sampling thereafter. On 5 dpi, animals (*n* = 4) from one group of each variant were euthanized and the respiratory tract (nasal turbinate, trachea and lungs) was sampled for quantification of viral and genomic load, as well as for histopathology and virus antigen expression. All remaining animals were sacrificed 26 dpi.

A single group of hamsters (*n* = 4) previously inoculated intranasally with 10^3^ TCID50 of Bavpat-1 (isolate BetaCoV/Munich/BavPat1/2020; European Virus Archive Global #026V 03883; kindly provided by Dr. C. Drosten) was challenged at 13 dpi by intranasal inoculation with 5 x 10^4^ PFU Omicron BA.1 in a volume of 100 μl. Throat swabs were taken at 1, 3 and 5 dpi and the animals were weighed daily. All animals of this group were sacrificed at 5 dpi.

### RNA extraction and genome quantification with RT-qPCR

All throat swabs, nasal washes and tissue samples were used for RNA extraction and following quantification of SARS-CoV-2 genome by RT-qPCR. RNA extraction was performed as described previously [13]. A RT-qPCR targeting the E gene of SARS-CoV-2 was used as previously reported [14]. Ct values were compared to a standard curve derived from a titrated 614G virus stock.

### Plaque assay

All throat swabs, nasal washes and tissue samples that were taken post mortem at 5 dpi were used for quantification of viral loads via a plaque assay on Calu-3 cells as described before [15] (Supplemental material). Briefly, ten-fold serial diluted samples were added to monolayers of Calu-3 cells. Cells were incubated with inoculums at 37°C for 4 hours, washed once with PBS and then overlayed with 1.2% Avicel (FMC biopolymers) in Opti-MEM I (1X) + GlutaMAX for two days. Cells were fixed in 4% formalin for 20 minutes, permeabilized in 70% ice-cold ethanol and washed in PBS. Cells were blocked in 3% BSA (bovine serum albumin; Sigma) in PBS and stained with rabbit anti-nucleocapsid (Sino biological; 1:2000) in PBS containing 0.1% BSA, washed thrice, and stained with donkey anti-rabbit Alexa Fluor 488 (Invitrogen; 1:4000) in PBS containing 0.1% BSA. Plates were washed and scanned on the Amersham Typhoon Biomolecular Imager (channel Cy2; resolution 25 μm; GE Healthcare). All staining steps were performed at room temperature for one hour. Plaque assay analysis was performed using ImageQuant TL 8.2 software (GE Healthcare).

### Histopathology and immunohistochemistry

Nasal turbinates, trachea and lungs were fixed in 4% neutral-buffered formalin, embedded in paraffin, and sectioned at 4 μm. Sections of all tissue samples were stained with hematoxylin and eosin for histopathological analysis, and consecutive sections were stained by immunohistochemistry for SARS-CoV-2 antigen expression, as described previously [16].

### Statistical analysis

All statistical analyses were performed using GraphPad Prism 9 software (La Jolla, CA, USA). Applied tests are indicated in according figure legends.

## Results

### Clinical manifestation and shedding after inoculation of Syrian golden hamsters with Omicron BA.1 or Delta

Following intranasal inoculation of Syrian golden hamsters with the Omicron BA.1 and Delta variants of SARS-CoV-2, differences in clinical manifestation were monitored by body weight measurements of individual animals (**Figure 1A**). Compared to mock-infected hamsters, infection with the Delta variant resulted in a significant body weight loss of maximum 14%, peaking at 4-5 dpi. No weight loss was observed in animals infected with the Omicron BA.1 variant and animals gained weight comparable to mock-infected animals.

**Figure 1.**
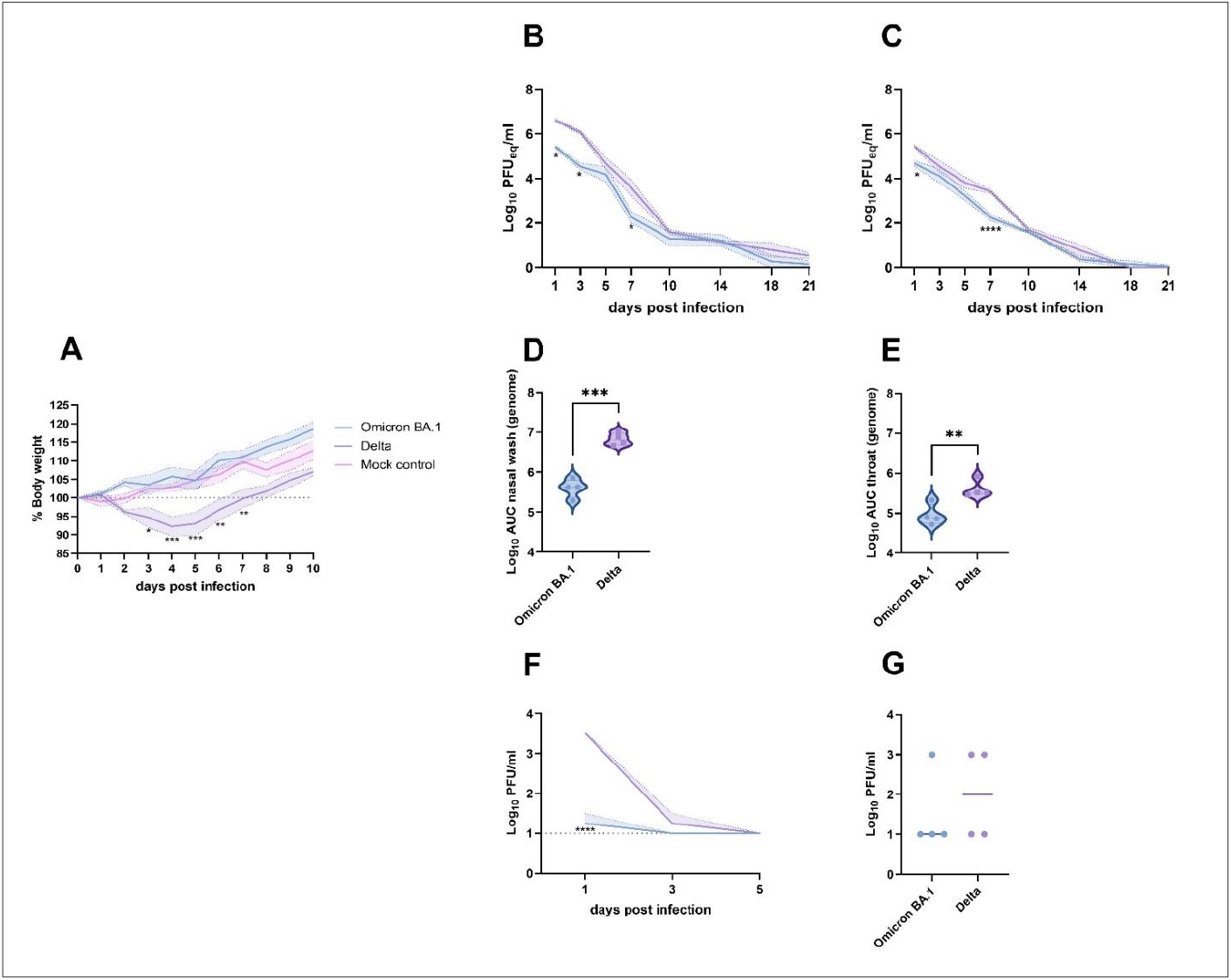
Clinical manifestation and shedding after Omicron BA.1 and Delta inoculation. (**A**) Relative body weight of hamsters inoculated with Omicron BA.1, Delta and mock-inoculated hamsters. The mean of each group is shown as solid line, SEM displayed as shading. Significant statistical differences between the three groups were calculated with a 2-way ANOVA with Tukey’s multiple comparison test. Significant differences to the mock control are indicated with * (*: p<0.05; **: p<0.01; ***: p<0.001). (**B,C**) Kinetics of respiratory shedding detected in (**B**) nasal washes and (**C**) throat swabs. The mean of each group is shown as solid line, SEM displayed as shading. Significant statistic differences (*: p<0.05; ****: p<0.0001) were calculated with 2-way ANOVA with Dunnett’s correction for multiple comparisons. (**D,E**) Cumulative shedding depicted as area under the curve (AUC) in (**D**) nasal washes and (**E**) throat swabs. Significant statistic differences were calculated with unpaired t-test (**: p<0.01; ***:p<0.001).(**F,G**) Detection of infectious virus. (**F**) Infectious virus in nasal washes. The mean of each group is shown as solid line, SEM displayed as shading. Significant statistic differences (****: p<0.0001) were calculated with 2-way ANOVA with Dunnett’s correction for multiple comparisons. (**G**) Infectious virus in throat swabs at 1 dpi. The mean of each group is depicted as horizontal line. Significant statistic differences were calculated with unpaired t-test.

To evaluate if Omicron BA.1 and Delta variants were shed to a varying extent via the upper respiratory tract, both nasal washes and throat swabs were taken regularly throughout the experiment. SARS-CoV-2 genome was detected in the nasal wash and throat swabs of all hamsters until 21 dpi (**Figure 1B,C**). Occasionally, significant differences in detection of genomic RNA between Omicron BA.1 and Delta were observed, that were confirmed by the comparison of cumulative shedding (**Figure 1D,E**). In nasal washes, infectious virus was only found up to 3 dpi (**Figure 1F**). Significantly lower amounts of infectious virus were detected in the nasal washes of Omicron BA.1 infected animals on 1 dpi, compared to animals infected with Delta (p<0.0001). Infectious virus in throat swabs was only detectable on 1 dpi, with no significant differences between the two groups (**Figure 1G**).

### Pathogenesis of Omicron BA.1 infection in the Syrian golden hamster

Despite the fact that Omicron BA.1 infected animals did not show weight loss, gross postmortem examination at 5 dpi revealed single or multiple foci of pulmonary consolidation, visible as well-delimited, slightly raised, dark red areas, covering 50-90% of the lung surface (**Figure 2A**). Due to animal-to-animal variation, no clear differences in pulmonary consolidation were observed between Delta and Omicron BA.1. The main histopathological changes in the lungs of both Omicron BA.1 and Delta infected animals were multiple areas of diffuse alveolar damage, characterized by flooding of alveolar lumina by variable proportions of neutrophils and macrophages, mixed with fibrin and oedema fluid; and by type II pneumocyte hyperplasia (**Figure 3**). While the histopathological features were qualitatively similar between hamster inoculated with the two strains, a significant difference in the percentage of inflamed lung area was observed, with an average of 78% of the lung area affected in Delta variant infected animals, compared to 38% in Omicron BA.1 infected animals (**Figure 2B**). By immunohistochemistry, mainly type I pneumocytes at the edges of the lesions expressed viral antigen with less than 2% of the lung area being positive. No significant difference in the amount of viral antigen expression was observed between Omicron BA.1 and Delta (**Figure 2C**).

**Figure 2.**
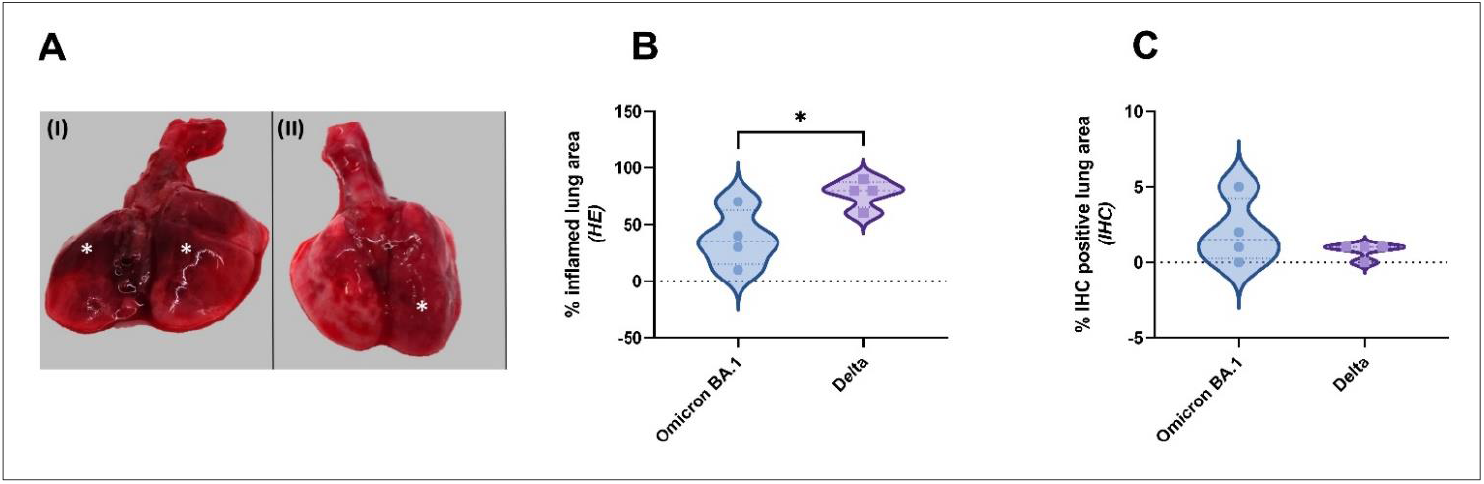
Gross pathological lesion and quantitative scoring of histopathological changes and viral antigen expression in Omicron BA.1 or Delta inoculated hamsters. (**A**) Representative gross pathological lesions of animals inoculated with (I) Omicron BA.1 or (II) Delta. Foci (*) of pulmonary consolidation. (**B**) Percentage of inflamed lung areas detected in HE staining and (**C**) % of viral antigen-positive lung areas detected in IHC. Significant statistical differences between two groups were calculated with unpaired t-test (*: p<0.05).

**Figure 3.**
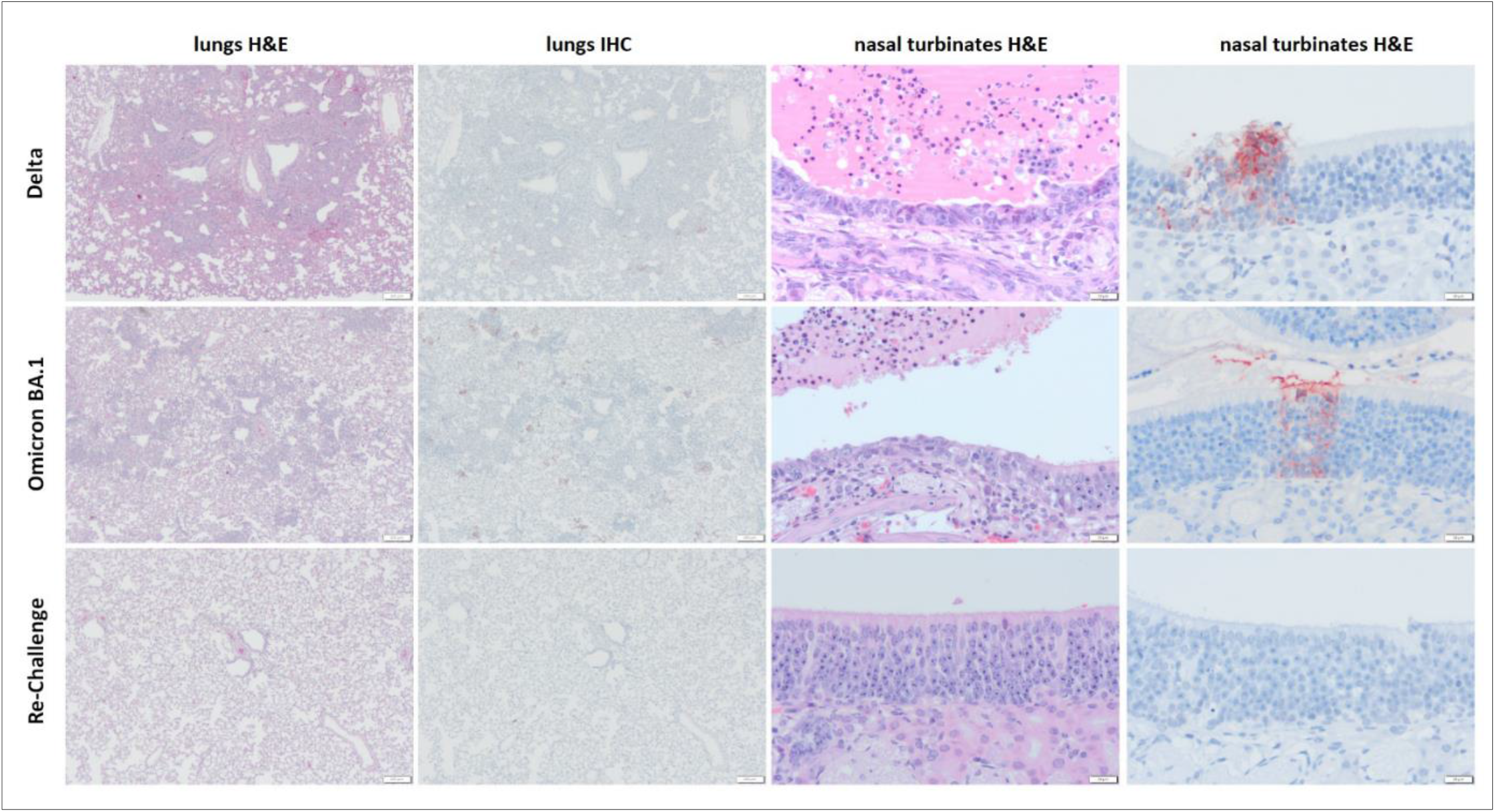
Histopathological changes and viral antigen expression in nasal turbinates and lungs of Omicron BA.1 and Delta inoculated hamsters. H&E staining and IHC of lungs and nasal turbinates of Syrian golden hamsters, infected with Delta or Omicron BA.1 or convalescent hamsters upon re-challenge with Omicron BA.1. (Re-Challenge). Bars indicate 200 μm for image of the lungs and 20 μm for the nasal turbinates.

The main histopathological changes in the nasal turbinates of both Omicron BA.1 and Delta infected animals were necrosis and inflammation of the olfactory mucosa. This was characterized by the presence of many neutrophils mixed with cellular debris and fibrin in the lumina of the nasal turbinates, as well as infiltration with variable numbers of neutrophils in the underlying olfactory mucosa. In addition, there was multifocal attenuation of the olfactory mucosa, indicative of previous loss of epithelial cells due to infection at earlier time points. Both the amount of neutrophil infiltration and of olfactory mucosal attenuation appeared to be increased in Delta-infected animals than in Omicron BA.1-infected animals. SARS-CoV-2 antigen was mainly found in cellular debris within the lumina of nasal turbinates, again indicative of damage at earlier timepoint. Additionally, small foci within the olfactory mucosa were found to be positive for viral antigen (**Figure 3**). Furthermore, respiratory epithelial cells were occasionally found to carry viral antigen in Delta, but not Omicron BA.1 infected animals. There were no clear differences in the amount of virus antigen expression between Omicron BA.1 - and Delta-infected animals. Neither virus antigen expression nor lesions were found in the tracheas of any of the animals.

### SARS-CoV-2 in respiratory tissues of Omicron BA.1 infected Syrian golden hamsters

Besides of pulmonary lesions and the presence of viral antigen, respiratory tissue was also evaluated for the presence of viral RNA via RT-qPCR and infectious virus. SARS-CoV-2 genome copies were lower in respiratory tissues of Omicron BA.1 inoculated hamsters, compared to hamsters infected with the Delta variant. However, statistically significant differences were only detected in the nasal turbinates when compared to Delta infected animals (**Figure 4A**). Infectious virus was mainly found in the nasal turbinates, although low virus titers were also found in the lungs of one out of four Omicron BA.1 and one out of four Delta infected hamsters (**Figure 4B**). No infectious virus was found in the trachea.

**Figure 4.**
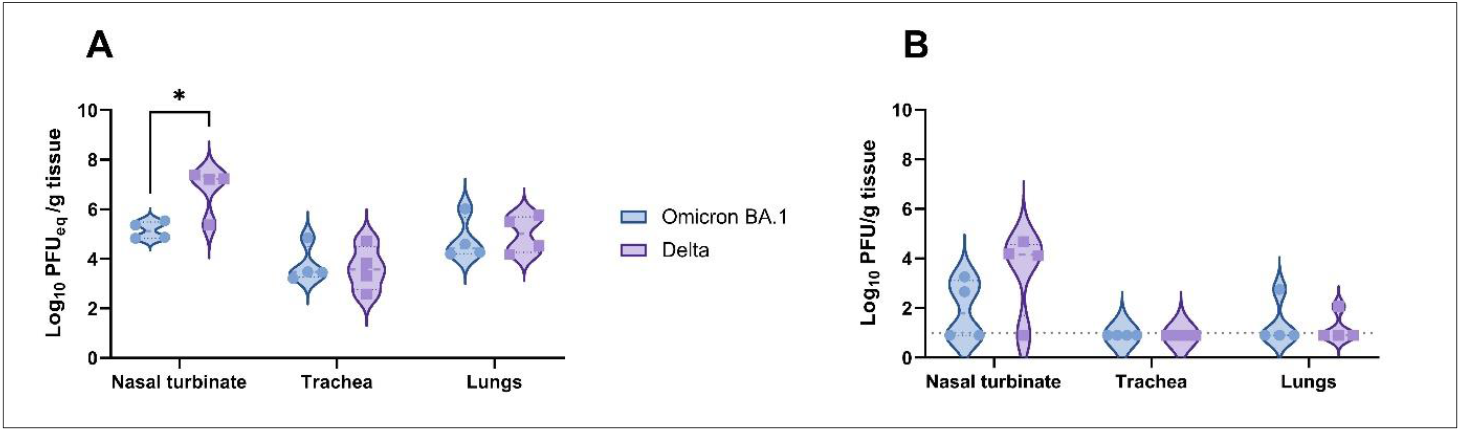
Detection of SARS-CoV-2 in respiratory tissue of Omicron BA.1 or Delta inoculated hamsters. (**A**) Detection of SARS-CoV-2 genome in respiratory tissue of inoculated hamsters. Significant statistical differences were calculated with unpaired t-test (*: p<0.05). (**B**) Detection of infectious virus in respiratory tissue of Omicron BA.1 and Delta infected Syrian golden hamsters. Significant statistical differences were calculated with unpaired t-test.

### Protective effect of previous inoculation with 614G SARS-CoV-2 on re-challenge with Omicron BA.1

In heterologous re-challenged animals, no changes in the percentage of body weight were observed (**Figure 5A**). At 3 dpi and 5 dpi, significantly less SARS-CoV-2 genome was detected in throat swabs of re-challenged animals (p<0.0001) compared to Omicron BA.1 infected hamsters (**Figure 5B**). No infectious virus was detected in any throat swabs of the re-challenged animals. No gross lesions were observed in the lungs of re-challenged animals (**Figure 5C**).

**Figure 5.**
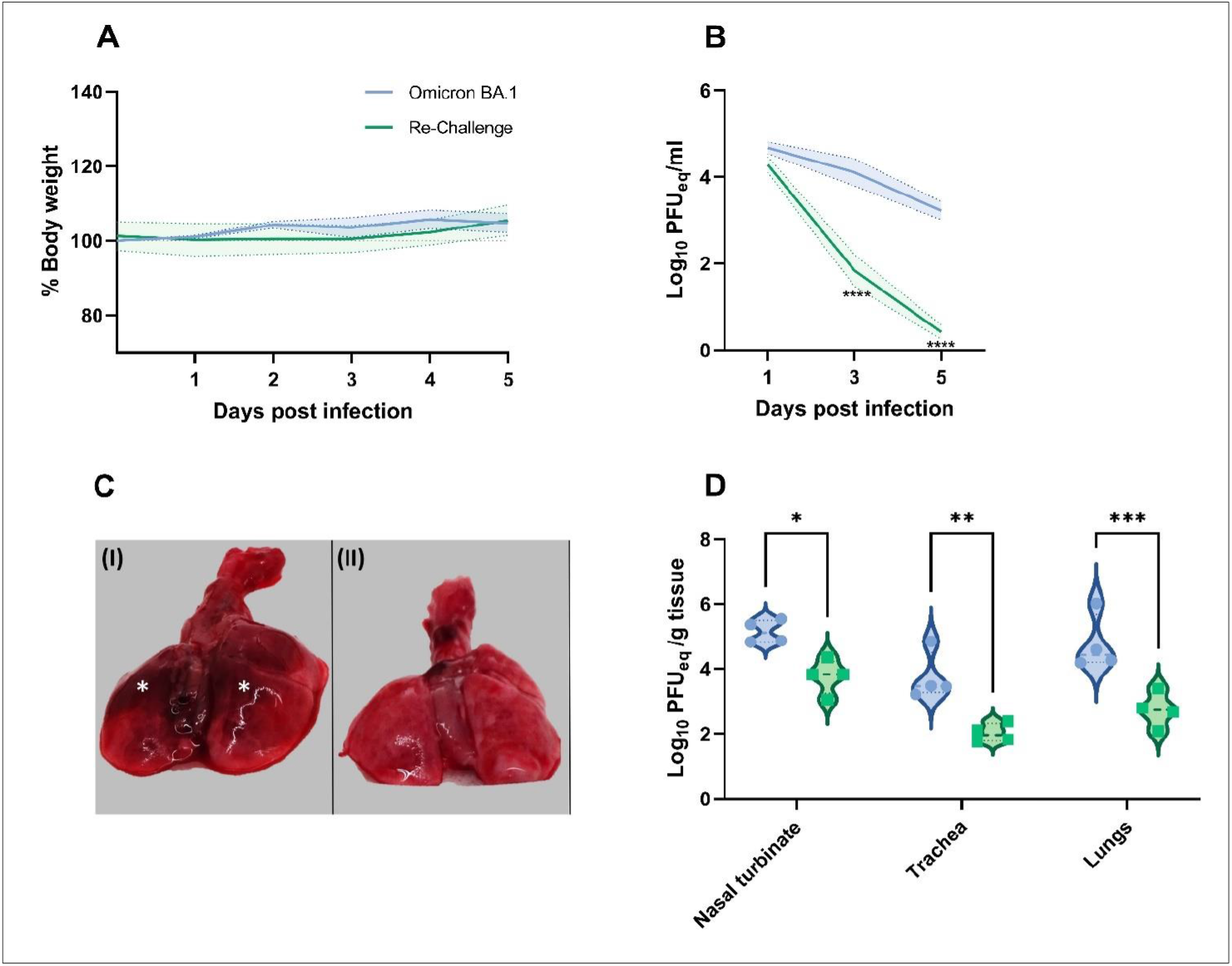
Effect of Omicron BA.1 re-infection in 614G convalescent Syrian golden hamsters. (**A**) Relative body weight of immunologically naïve hamsters infected with Omicron and convalescent hamsters re-challenged with Omicron BA.1. The mean of each group is shown as solid line, SEM displayed as shading. Significant statistic differences were calculated with 2-way ANOVA with Dunnett’s correction for multiple comparisons. (**B**) Detection of viral genome in throat swabs of immunologically naïve hamsters infected with Omicron BA.1 and convalescent hamsters re-challenged with Omicron. The mean of each group is shown as solid line, SEM displayed as shading. Significant statistic differences (****: p<0.0001) were calculated with 2-way ANOVA with Dunnett’s correction for multiple comparisons. (**C**) Representative gross pathological lesions of animals inoculated with (I) Omicron BA.1^1^ or (II) 614G convalescent and Omicron BA.1 re-challenged animals. Foci (*) of pulmonary consolidation. (**D**) Detection of SARS-CoV-2 genome^2^ in respiratory tissue of infected hamsters. Significant statistic differences were calculated with unpaired t-test (*: p<0.05). ^1,2^ Data shown for comparison to Omicron BA.1 inoculated hamsters are based on data shown in Figure 1 (^1^) and 4 (^2^).

The genomic viral burden in nasal turbinates, trachea and lungs was significantly higher in naïve hamsters infected with Omicron BA.1, compared to re-challenged animals. This difference was most pronounced in the lungs (p=0.0004) (**Figure 5D**). No infectious virus was found in any tissue of re-challenged animals. No histopathological changes or viral antigen expression were found in the lungs or nasal turbinates of re-challenged hamsters (**Figure 3**).

## Discussion

Following the identification of SARS-CoV-2 as the cause of coronavirus disease-2019 (COVID-19), the virus has spread rapidly throughout the world. Since its emergence, viral genomic sequence analysis has allowed tracking of the emergence of SARS-CoV-2 variants. While many of the observed mutations in the SARS-CoV-2 genome are neutral, or deleterious, some mutations are associated with an increase of viral fitness. These changes may include immune escape as well as changes in pathogenicity, infectivity and transmissibility [3]. This study shows that in the hamster model, the recently emerged VOC Omicron BA.1, does not cause body weight loss but still induces gross- and histopathological changes in the upper and lower respiratory tract with decreased viral shedding compared to the Delta variant of SARS-CoV-2. However, a previous inoculation with the ancestral 614G SARS-CoV-2 confers complete protection from pneumonia or viral replication in the respiratory tract. As shown by us and others, inoculation of hamsters with ancestral SARS-CoV-2 614G results in significant but transient body weight loss peaking at 4-5 dpi in the absence of overt respiratory signs, making body weight loss a valuable readout for clinical impact of SARS-CoV-2 inoculation [17]. Here, we show that the Delta variant causes significant body weight loss in hamsters, whereas no weight loss can be observed after infection with Omicron BA.1 in a direct comparison. Although most other studies observed a limited body weight loss or at least decreased weight gain compared to non-infected animals, a marked reduction of disease severity after infection with Omicron BA.1 is reported throughout all previous studies [10, 11], similar to what is observed in humans [18].

Previous studies in clinical settings have shown that viral genome of certain SARS-CoV-2 variants is shed at higher levels than for other variants. This increased shedding in part has been suggested to contribute to in the increased spread of these variants [19]. However, recent analysis of human specimens are indicative for no major differences in shedding between the SARS-CoV-2 variants Delta and Omicron BA.1 [20]. We demonstrated that the kinetics of viral genome shedding were not significantly different between Delta and Omicron BA.1 infected hamsters. Different to previous reports of other groups, where Omicron BA.1 RNA was only shed until 9 dpi and Delta until 12 dpi [21], we observed shedding until 21 dpi. Recent studies also demonstrated that the transmissibility in a non-contact transmission setup is increased for Omicron infected animals compared to Delta infected animals. Both groups had comparable titers in nasal turbinates of index animals with lower viral shedding in Omicron BA.1 infected animals during the course of infection [21]. Collectively, this indicates that evaluation of viral shedding in the hamster model cannot solely elucidate the transmissibility of different VOCs.

While no body weight loss was observed in Omicron BA.1 infected animals, these animals still showed diffuse alveolar damage and viral antigen expression, although milder compared to infection with Delta and to previously reported inoculation with the ancestral 614G strain [22]. This observation stands in contrast to previous reports of Omicron BA.1 infections in hamsters, describing limited or no inflammation in the lower respiratory tract [10, 11, 21]. A lower inoculation dose of 10^3^ PFU in 50 μl, compared to 5.0 x 10^4^ PFU in 100 μl used in this study, may explain the differences for one previous study [3]. However, two other studies used even higher doses of 10^5^ PFU in 100 μl [12] or 10^5^ PFU in 30 μl [4] and only saw limited pathology. In contrast to these studies, where viral stocks were propagated on Vero cells, we used viruses grown on Calu-3 cells, shown to have a cleaved spike protein [23].

While the observed reduction in disease severity in humans may (in part) be the result of intrinsic properties of the virus, the presence of pre-existing immunity, due to vaccination or previous infection, may also affect the disease outcome. We previously showed that the Omicron BA.1 variant is antigenically distinct from the other SARS-CoV-2 variants [24], and several studies have shown that neutralizing monoclonal antibodies against the ancestral SARS-CoV-2 strain have limited neutralizing capacity against the Omicron BA.1. Interestingly, despite the antigenic distance and early time point for re-infection, the 614G convalescent group was completely protected from any pathological alterations of the respiratory tract after infection with Omicron BA.1. This indicates a strong cross-protective effect of heterologous re-infections and will need further investigations of the mechanistic background and scope of this cross-protection in prospective studies. Additionally, the experimental setup of re-challenge of convalescent animals can be a useful tool to study if observed decrease of disease severity is mainly driven by viral factors or pre-existing immunity.

In conclusion, with the continuing emergence of SARS-CoV-2 variants, models to evaluate potential phenotypic changes associated with mutations are invaluable in assessing the impact on public health. In hamsters, the currently circulating dominant SARS-CoV-2 Omicron BA.1 variant, shows reduced viral shedding and disease, recapitulating the phenotype seen in human cases. Interestingly, remaining pathogenicity in the lower respiratory tract can still be detected without being reflected in clinical manifestation. In summary, we show that the hamster model remains useful in assessing the impact of SARS-CoV-2 variants on disease severity and the role of pre-existing immunity. Thus, the model continues to be an important tool in the evaluation of pathogenicity of different SARS-CoV-2 variants and studying the efficacy of intervention strategies.

## Acknowledgments

We thank J.M. Fentener van Vlissingen, Ingeborg van Middelkoop, Rianne Stam, Vincent Duiverman and Vincent Vaes for assistance with the animal studies.

## Author contributions

Conceptualization, B.R., M.R., R.F. B.H.; investigation, M.R., D.N., D.v.R., K.S.S., R.D.d.V, P.v.R., T.K. and B.R.; resources, M.L., C.G.v.K., M.P.G.K.; supervision, B.R., R.D.d.V and B.H.; writing, original draft, B.R., M.R., and B.H.; writing–review and editing, all authors; funding acquisition: M.P.G.K., R.F. B.H., B.R.,

## Data and materials availability

Viruses must be obtained through an MTA.

