## Supplemental material for "Pulmonary lesions following inoculation with the SARS-CoV-2 Omicron BA.1 (B.1.1.529) variant in Syrian golden hamsters"

**Supplemental materials**

**Animals and Ethical Statement**

Research involving animals was conducted in compliance with the Dutch legislation for the protection of animals used for scientific purposes (2014, implementing EU Directive 2010/63) and other relevant regulations. The licensed establishment where this research was conducted (Erasmus MC) has an approved OLAW Assurance # A5051-01. Research was conducted under a project license from the Dutch competent authority and the study protocol (#17-4312) was approved by the institutional Animal Welfare Body. Animals were housed in groups of 2 animals in filter top cages (T3, Techniplast), in Class III isolators allowing social interactions, under controlled conditions of humidity, temperature and light (12-hour light/12-hour dark cycles). Food and water were available ad libitum. Animals were cared for and monitored (pre- and post-infection) daily by qualified personnel. All animals were allowed to acclimatize to husbandry for at least 7 days. For unbiased experiments, all animals were randomly assigned to experimental groups. The animals were anesthetized (3-5% isoflurane) for all invasive procedures. Hamsters were euthanized by cardiac puncture under isoflurane anesthesia and cervical dislocation.
